# Flv1-4 proteins function in versatile combinations in O_2_ photoreduction in cyanobacteria

**DOI:** 10.1101/561456

**Authors:** Anita Santana-Sánchez, Daniel Solymosi, Henna Mustila, Luca Bersanini, Eva-Mari Aro, Yagut Allahverdiyeva

## Abstract

Flavodiiron proteins (FDPs) constitute a group of modular enzymes widespread in all life Domains. *Synechocystis* sp. PCC 6803 has four FDPs (Flv1-4) essential for photoprotection of photosynthesis. A direct comparison of the Mehler-like reaction (O_2_ photoreduction) in high Ci (3% CO_2_, HC) and low Ci (air level CO_2_, LC) acclimated cells demonstrated that the Flv1/Flv3 heterodimer is responsible for an efficient steady-state O_2_ photoreduction under HC, with *flv2* and *flv4* expression strongly down-regulated. Conversely, under LC conditions Flv1/Flv3 acts only as a transient electron sink due to competing withdrawal of electrons by the highly induced NDH-1 complex. Further, *in vivo* evidence is provided indicating that Flv2/Flv4 contributes to the Mehler-like reaction when naturally expressed under LC conditions, or when artificially overexpressed under HC. The O_2_ photoreduction driven by Flv2/Flv4 occurs down-stream of PSI in a coordinated manner with Flv1/Flv3 and supports slow and steady-state O_2_ photoreduction.

## Introduction

A-type flavodiiron proteins (Flvs or FDPs) were originally identified in strict and facultative anaerobes among Bacteria, Archaea and Protozoa and were considered to function in O_2_ and/or NO detoxification (Wasserfallen et al., 1998; Gonçalves et al., 2011; Folgosa et al., 2018). All FDPs share two conserved structural domains: the N-terminal metallo-*β*-lactamase-like domain, harboring a non-heme diiron center, where O_2_ and/or NO reduction takes place; and the C-terminal flavodoxin-like domain, containing a flavin mononucleotide (FMN) moiety. The structures of FDPs in anaerobic prokaryotes and eukaryotic protozoa have been resolved as homooligomers (dimer or tetramer comprised of two dimers) arranged in a “head-to-tail” configuration, so that the diiron center of one monomer and the FMN of the other monomer are in close proximity to each other, which ensures rapid electron transfer between the two cofactors.

C-type FDPs, specific to oxygenic photosynthetic organisms, hold an additional flavin-reductase-like domain, coupled with extra cofactors (Romão et al., 2016; Folgosa et al., 2018). *Synechocystis* sp. PCC 6803 (hereafter, *Synechocystis*) possesses four genes encoding FDPs: *sll1521* (Flv1), *sll0219* (Flv2), *sll0550* (Flv3) and *sll0217* (Flv4). Recently resolved crystal structure of truncated Flv1 from *Synechocystis* revealed a monomeric form with a ‘bent’ configuration (Borges et al. 2019). Photosynthetic FDPs gained attention first in 2002 when recombinant *Synechocystis* Flv3 protein was shown to function in O_2_ reduction to water without producing ROS (Vicente et al., 2002). Later it was demonstrated that *Synechocystis* Flv1 and Flv3 proteins function *in vivo* in the photoreduction of O_2_ downstream to PSI (Helman et al., 2003). Since then, extensive research has been pursued to reveal the crucial function of Flv1 and Flv3 (and their homologs, FLVA and FLVB in other photosynthetic organisms) as a powerful sink of excess photosynthetic electrons which safeguards PSI and secures the survival of oxygenic photosynthetic organisms under fluctuating light (Allahverdiyeva et al., 2013; Gerotto et al., 2016; Chaux et al., 2017; Jokel et al., 2018) or under the repetitive short-saturation pulses (Shimakawa et al., 2017). The Flv1- and Flv3-mediated alternative electron transport to O_2_ was named as the Mehler-like reaction, being a widespread pathway, operating in all photosynthetic organisms from cyanobacteria up to gymnosperms, but lost in angiosperms (Allahverdiyeva et al., 2015; Ilík et al., 2017).

The Flv2 and Flv4 proteins are encoded by an operon together with a small membrane protein, Sll0218. The *flv4-sll0218-flv2* (hereafter *flv4-2*) operon is strongly induced in low Ci (atmospheric 0.04% CO_2_ in air, LC) and high light conditions (Zhang et al., 2009). The operon structure is highly conserved in the genome of many *β*-cyanobacteria (Zhang et al., 2012; Bersanini et al., 2014). The *flv4-2* operon-encoded proteins have been reported to function in photoprotection of Photosystem (PS) II, by acting as an electron sink presumably transporting electrons from PSII or plastoquinone (PQ) pool to an unknown acceptor (Zhang et al., 2009, 2012; Bersanini et al., 2014; Chukhutsina et al., 2015). Since *flv2*, *sll0218* and *flv4* are co-transcribed, the contribution of each single protein of the operon to PSII photoprotection has been difficult to dissect. Recent data examining distinct and specific roles of the Flv2/Flv4 heterodimer and the Sll0218 protein (using a set of different mutants deficient only in Sll0218 or in Flv2 and Flv4) demonstrated that the majority of observed PSII phenotypes were actually due to the absence of Sll0218, thus leading to the conclusion that Sll0218 contributes to PSII repair and stability (Bersanini et al., 2017). However, the exact donor and acceptor of the Flv2 and Flv4 proteins *in vivo* have not been identified and possible cross-talks between all four FDPs have not been established so far, thus limiting our understanding of the function of FDPs on cellular level.

In order to shed light on *in vivo* function of Flv2 and Flv4 and to untangle the function of Flv1/Flv3 from the one of Flv2/Flv4 heterooligomer, we used here a specific set of FDP mutants. The set included: (i) the Δ*flv1*/Δ*flv3* mutant deficient in both Flv1 and Flv3 proteins (Allahverdiyeva et al., 2011); (ii) the Δ*flv2* mutant, which does not express the Flv2 protein, but retains a low amount of Flv4 and the WT level of Sll0218 (Zhang et al., 2012); (iii) the Δ*flv4* mutant which is deficient in the accumulation of three *flv4-2* operon proteins (Zhang et al., 2012); (iv) Δ*sll0218*, which lacks the small Sll0218 protein, but expresses the Flv2 and Flv4 proteins (Bersanini et al., 2017); (v) Δ*flv3*/Δ*flv4*, which represents a deficiency in all four FDPs, whereby the absence of Flv3 results in a strong decrease in Flv1 (Mustila et al., 2016) and the inactivation of Δ*flv4* affects the expression of the whole *flv4*-*2* operon (Zhang et al., 2012); and finally (vi) the *flv4-2* operon overexpression strain, *flv4-2*/OE, expressing high amounts of Flv2, Flv4 and Sll0218 (Bersanini et al., 2014).

Here, we provide *in vivo* evidence for Flv2/Flv4 mediated O_2_ photoreduction in *Synechocystis*. Unlike the rapidly responding powerful Flv1 and Flv3, the more slowly functioning Flv2 and Flv4 proteins are dispensable for survival under fluctuating light intensities. The expression of *flv4* and *flv2* under LC was found to be regulated by pH of the growth media, with significant downregulation observed under strong alkaline pH conditions. Results from this study provide important insights into the response of photosynthetic organisms to changes in Ci and into how they regulate the availability of electron sinks.

## Results

### 1. Extent and kinetics of the Mehler-like reaction in cells acclimated to low (LC) and high Ci (HC) conditions

Application of membrane inlet mass spectrometry (MIMS) with ^18^O-enriched oxygen allows differentiation between photosynthetic gross O_2_ production and O_2_ uptake under illumination. The *flv4-2*/OE cells, accumulating high amounts of Flv2, Sll0218 and Flv4 both in LC and high Ci (> 1% CO_2_ in air, HC) conditions (Bersanini et al., 2014), demonstrated substantially higher O_2_ photoreduction rates compared to the respective WT (Figure 1A, 1B, 1D). The Flv3 protein level was similar in *flv4-2*/OE and wild-type (WT) cells grown both at LC and HC (Figure 1C) thus strongly supporting the contribution of the *flv4-2* operon proteins to O_2_ photoreduction *in vivo* during illumination.

**Figure 1.**
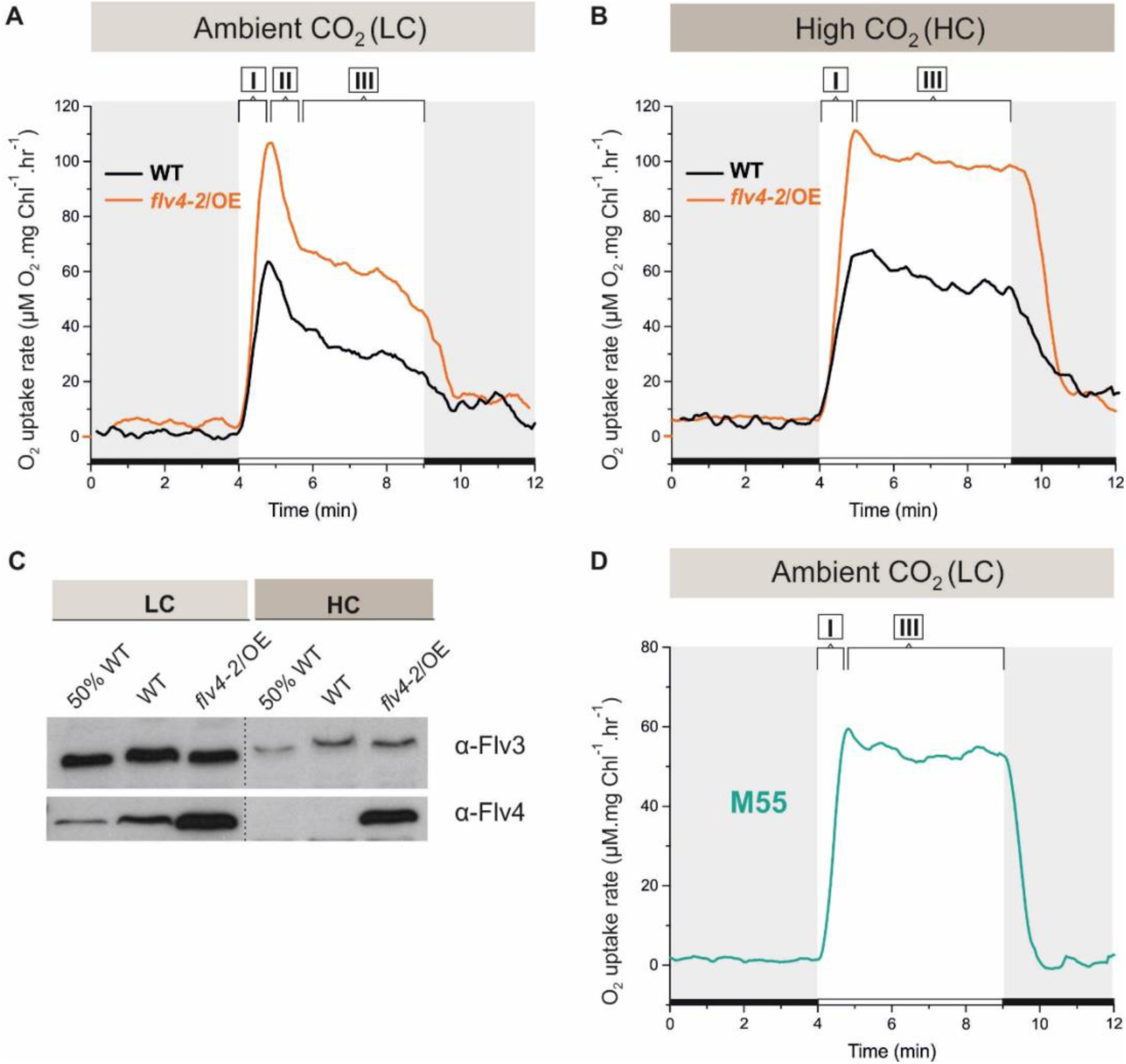
O_2_ reduction rates and Flv3 and Flv4 protein accumulation in cells grown in low (LC) and high CO_2_ (HC). (A, B) O_2_ reduction rate of WT, *flv4-2*/OE and (D) the M55 mutant (Δ*ndhB)* was recorded in darkness (grey background) and under illumination (white background). The experiment was conducted in 3 independent biological replicates and a representative plot is shown. (Figure 1-source data 1). (C) Immunoblot detection of Flv3 and Flv4 in WT and *flv4-2*/OE. Pre-cultures were grown in BG-11, pH 8.2 under 3% CO_2_ (HC) for 3 days, after that cells were harvested and resuspended in fresh BG-11, pH 8.2 at OD_750_=0.2. The experimental cultures were grown under HC or under LC. For the MIMS experiments the cells were harvested and resuspended in fresh BG-11, pH 8.2 at 10 µg Chl *a* mL^−1^. O_2_ photoreduction was recorded during the transition from darkness to high-light intensity of 500 µmol photons m^−2^s^−1^. Different phases of O_2_ photoreduction kinetics are indicated as {I}, {II}, {III}.

As reported earlier, the Ci level has a remarkable effect on the expression of FDPs at transcript and protein level: Flv2, Flv4 and Flv3 have been strongly upregulated under LC (Zhang et al., 2009; Wang et al., 2004; Battchikova et al., 2010), and down-regulated upon a shift to HC (Zhang et al., 2009; Hackenberg et al., 2012; Figure 1C). Nevertheless, a direct comparison of the efficiency and kinetics of the Mehler-like reaction in HC- and LC-acclimated cells has not been reported, thus making the contribution of different FDPs to O_2_ photoreduction difficult to assess. Therefore, as an initial experiment, we evaluated the activity of the Mehler-like reaction in *Synechocystis* cells grown at LC and HC (3% CO_2_) conditions, at pH 8.2.

The following source data and figure supplements are available for Figure 1:

**Source data 1.** O_2_ reduction rates of WT, flv4-2/OE and the M55 mutant grown under different CO_2_ levels.

**Figure supplement 1.** *O_2_ reduction rates under high CO_2_.*

After a shift from darkness, WT cells demonstrated rapid induction of light-induced O_2_ uptake both at LC and at HC conditions (59 ± 6.4 and 56 ± 6.4 µmol O_2_ mg Chl *a* ^−1^ h^−1^, respectively). This fast induction phase is designated as {I} in Figure 1A and 1B. Yet, the kinetics of O_2_ photoreduction in LC-grown cells differed from those grown under HC. In the LC-grown WT cells, the fast induction phase {I} was followed by a clear biphasic quenching of O_2_ reduction, namely by the strong decay phase {II}, which continued about a minute, followed by a quasi-stable state, phase {III} (∼33 ± 5.9 µmol O_2_ mg Chl *a* ^−1^ h^−1^) during illumination. On the contrary, in HC-grown WT cell, the light-induced O_2_ reduction rate achieved in the phase {I} declined only slightly during the first 2-3 minutes (from ∼ 56 ± 7.7 to ∼ 48 ± 6.3 µmol O_2_ mg Chl *a* ^−1^ h^−1^). Thereafter it remained nearly at the same rate for at least 5 min (Figure 1B) of illumination. In *flv4-2*/OE cells, grown both in LC- and HC, light-induced O_2_ reduction was stronger that in WT. Nevertheless, the kinetic phases of O_2_ photoreduction in *flv4-2*/OE cells resembled those of respective WT cells, being relatively stable at HC and demonstrating a strong biphasic quenching at LC.

The Δ*flv2* and Δ*flv4* mutants grown under HC conditions demonstrated a similar O_2_ photoreduction pattern as WT, whereas the Δ*flv3*/Δ*flv4* and Δ*flv1*/Δ*flv3* mutants showed hardly any light-induced O_2_ reduction (Figure 1-Figure supplement 1). A negligible amount of Flv2 and Flv4 proteins in the WT cells grown at HC (Zhang et al., 2009, 2012; Figure 1C) explains the lack of contribution of these proteins to the Mehler-like reaction and confirms that the Flv1 and Flv3 proteins are responsible for the Mehler-like reaction under HC condition (Helman et al., 2003).

To disclose the reason for the fast decay of O_2_ photoreduction observed under LC conditions (Figure 1A), we first tested the putative competition between the NAD(P)H:quinone oxidoreductase (NDH-1) complex and FDPs for available photosynthetic electrons. The NDH-1 complex is a powerful machinery utilizing electrons for cyclic electron transport (CET) around PSI, CO_2_ uptake and respiration under LC conditions (Zhang et al., 2004, Schuller et al.., 2019). To this end, the O_2_ photoreduction was measured in the M55 mutant (Δ*ndhB*) deficient in the hydrophobic NdhB subunit (Ogawa et al., 1991) and thus lacking all NDH-1 complexes (Zhang et al., 2004). The M55 mutant demonstrated steady-state O_2_ photoreduction kinetics during the dark-to-light transition (Figure 1D), resembling in this respect the HC-grown WT cells (Figure 1B; Figure 1-Figure supplement 1) and thus confirming the possible competition for electrons between the NDH-1 complexes and FDPs under LC conditions.

### 2. Extent and kinetics of the Mehler-like reaction is strongly dependent on pH and the carbonate concentration of the growth medium

Next, the pH and the presence of carbonate in the growth medium were addressed as possible modulators of the extent and kinetics of the Mehler-like reaction and the accumulation of FDPs under LC conditions.

#### The effect of pH

The WT cells grown at pH 9 demonstrated a strong but only transient Mehler-like reaction: the O_2_ photoreduction rate reached its maximum during the first 30 s upon illumination, then quickly declined to the initial level of dark O_2_ uptake rate within 1 min of illumination (Figure 2, right panel). Similar to WT, the Δ*flv4* mutant cells demonstrated only a transient O_2_ photoreduction upon illumination. No significant O_2_ photoreduction could be recorded in the Δ*flv1*/Δ*flv3* and Δ*flv3*/Δ*flv4* mutants grown at pH 9.

**Figure 2.**
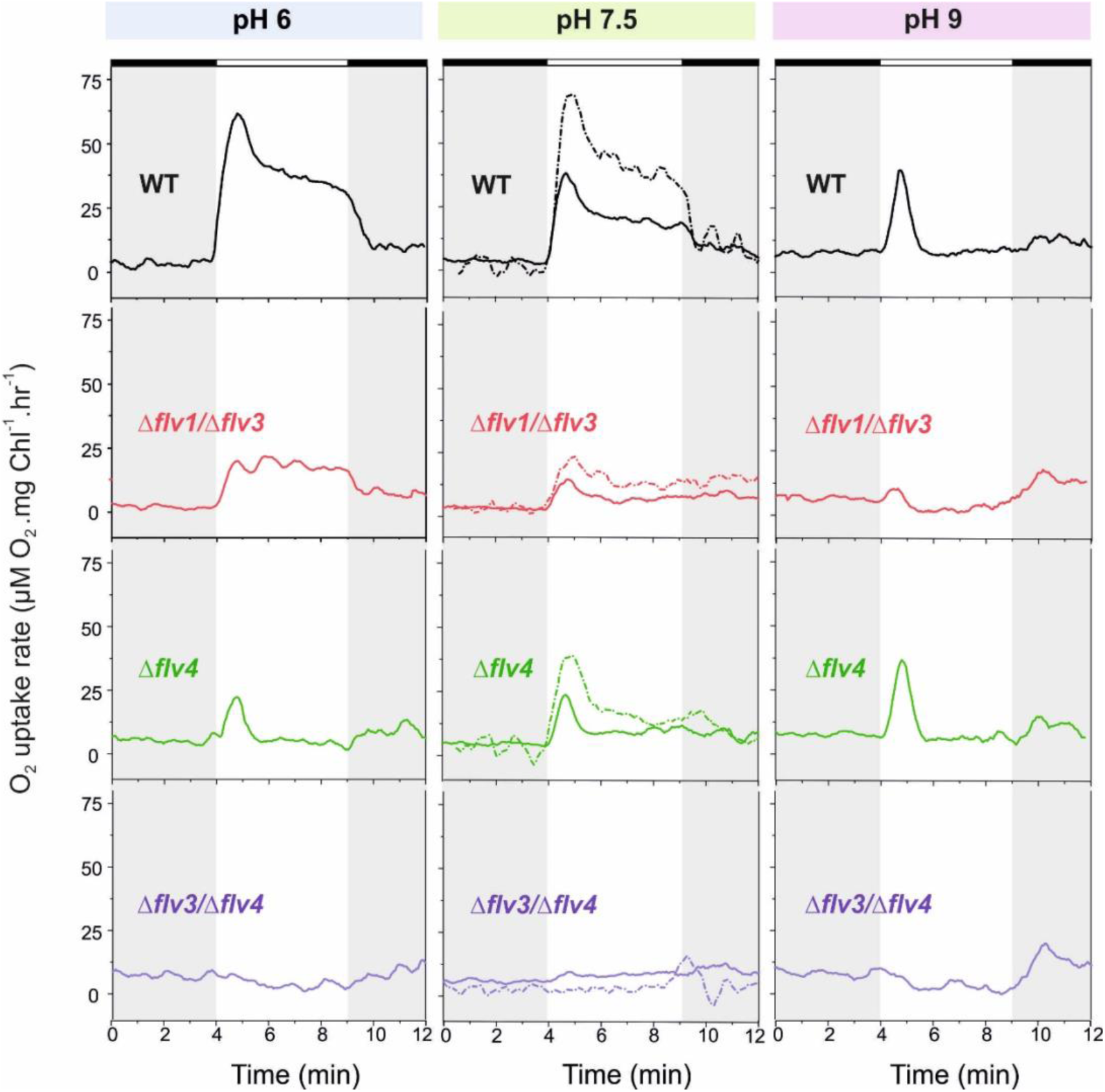
O_2_ reduction rates of WT and FDP mutants grown at different pH levels. O_2_ reduction rate was recorded in darkness (grey background) and under illumination with actinic white light at an intensity of 500 µmol photons m^−2^ s^−1^ (white background). Pre-cultures were grown under HC for 3 days at different pH levels. For MIMS experiments, cells were shifted to LC at OD750≈0.2 (same pH) and grown for 4 days. Exceptions were: (i) pH 6 experimental cultures were inoculated from pH 8.2 pre-cultures; and (ii) pH 7.5 pre-culture was shifted to LC in BG11, both with and without sodium carbonate (dotted line ‘-Na_2_CO_3_’) The experiment was conducted in 3 independent biological replicates (except experiment at pH 6 with n= 2 independent biological replicates) and a representative plot is shown. (Figure 2-Source data 1). In order to create comparable conditions for MIMS measurements, all cells were supplemented with 1.5 mM NaHCO_3_ prior to the measurements. Independent experiments performed on WT cells grown in the absence of Na_2_CO_3_, but supplied with 1.5 mM NaHCO_3_ prior to MIMS measurement showed no significant difference in O_2_ photoreduction rates (Figure 2-Figure supplement 1).

The following source data and figure supplement are available for Figure 2:

**Source data 1.** O_2_ reduction rates of WT and FDP mutants grown at different pH levels.

**Figure supplement 1.** *O_2_ photoreduction rates during the dark-to-light transition of WT cells with and without addition of 1.5 mM NaHCO_3_ prior MIMS measurements.*

***Figure supplement 2.** O_2_ photoreduction rates of the* Δ*flv2 and* Δ*sll0218 mutants grown at LC pH 7.5* and 8.2 *with and without Na_2_CO_3_*.

Immunoblotting using specific antibodies showed that, similarly to the WT cells grown under HC (Figure 1D), the Flv2 and Flv4 proteins were almost undetectable in WT grown under LC at pH 9 (Figure. 3A).

In line with protein data, the transcript levels of both *flv2* and *flv4* were significantly down-regulated in the cells grown at pH 9 (Figure 3B), suggesting a pH-dependent transcriptional regulation of the *flv4* and *flv2*. This is consistent with earlier transcriptional profiling experiments reporting downregulation of *flv2* and *flv4* transcripts after transferring *Synechocystis* from pH 7.5 to pH 10 (Summerfield and Sherman, 2008). Importantly, the accumulation of Flv3 was not affected at pH 9. These results strongly suggest that the conspicuous but transient O_2_ photoreduction observed in the WT and Δ*flv4* mutant cells at pH 9 originates mainly from the activity of Flv1/Flv3 heterodimer.

**Figure 3.**
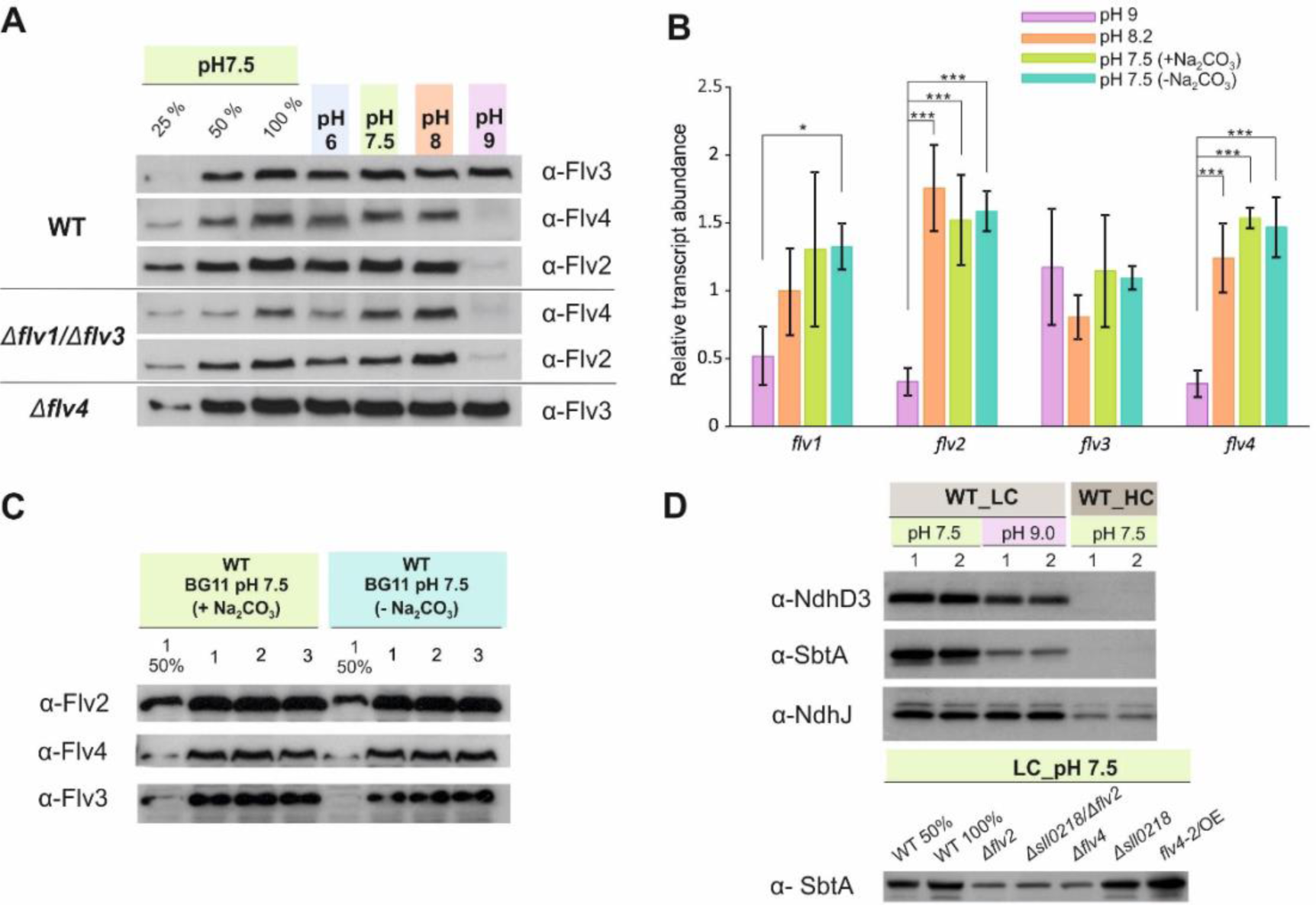
The effect of the pH of growth medium on the protein and transcript accumulation. (A, B) The effect of the pH and (B, C) absence of carbonate in the growth medium (A, C) on the protein and (B) transcript levels of FDP. (D) Protein immunoblots demonstrating the accumulation of bicarbonate transporter (SbtA) and NDH-1 subunits (NdhD3 and NdhJ) in the cells grown at different pH and CO_2_ concentration. Cells were pre-grown at different pH levels (+Na_2_CO_3_) under HC for 3 days, harvested, resuspended in fresh BG-11 (pH maintained), adjusted to OD_750_≈0.2 and shifted to LC for 4 days. At pH 7.5, the cells were grown at LC in the presence (+ Na_2_CO_3_) or in the absence (− Na_2_CO_3_) of sodium carbonate (B, C). Transcript abundance is presented as mean ± SD, n = 2-4 biological replicates, asterisks indicate a statistically significant difference to the WT (*P < 0.05; ***P<0.001) (Figure 3-source data 1). Numbers 1-3 indicate different biological replicates.

The WT cells grown at pH 6, at pH 7.5 (Figure 2, left and middle panels, respectively) and at pH 8.2 (Figure 1A) demonstrated a rapid induction of O_2_ reduction (phase {I}) followed by biphasic decay during illumination: a fast decay phase (phase {II}) and a quasi-stable phase (phase {III}) (Figure 1A and 2). The highest O_2_ photoreduction rate was observed in the WT cells grown at pH 6 (Figure 2).

Importantly, the Δ*flv1*/Δ*flv3* mutant also showed residual O_2_ photoreduction: only a small O_2_ uptake was noticeable at pH 7.5, whereas at pH 6 the O_2_ photoreduction rate was substantial and constant during 5 min of illumination of Δ*flv1*/Δ*flv3* (Figure 2). Unlike the Δ*flv1*/Δ*flv3* mutant, both the Δ*flv2* (Figure 2-Figure supplement 2) and Δ*flv4* (Figure 2) mutants showed a strong transient O_2_ photoreduction phase, peaking around the first 30 s of illumination and quickly decaying. This occurred at all tested pH levels. These results together with the ones that demonstrated highly increased rates of O_2_ photoreduction in the overexpression *flv4-2*/OE strain (Figure 1B) collectively confirmed the *in vivo* involvement of both the Flv2 and Flv4 proteins in O_2_ photoreduction. The O_2_ photoreduction kinetics of the Δ*sll0218* mutant resembled that of the WT (Figure 1-Figure supplement 1 and Figure 2-Figure supplement 2), thus implying that the Sll0218 protein does not contribute to the Mehler-like reaction under the HC and LC conditions studied here. These results made us to exclude the Δ*sll0218* mutant from any further experiments presented in this section.

The data presented above allowed making preliminary conclusions to be made about the origin of the different kinetic phases of O_2_ photoreduction. Since a transient O_2_ photoreduction was characteristic for the WT, Δ*flv2* and Δ*flv4* cells, but hardly detectable in Δ*flv1*/Δ*flv3*, it is conceivable that the Flv1/Flv3 heterodimer is mostly responsible for the strong and transient O_2_ uptake during dark-light transitions, whilst Flv2/Flv4 contribute to the steady-state O_2_ photoreduction at LC (see Δ*flv1*/Δ*flv3* particularly at pH 6, Figure. 2). The complete lack of O_2_ photoreduction in the Δ*flv3*/Δ*flv4* mutant (representing deficiency of all four FDPs) is in line with this hypothesis.

Yet the overall complexity of obtained results called for further investigation. Since not only the FDPs but also distinct variants of the NDH-1 complex as well as HCO_3_^−^ transporters (Zhang et al, 2004) are known to respond to the CO_2_ and pH levels of the growth medium, we next evaluated by immunoblotting the abundances of NdhD3, representing a low Ci inducible NDH-1MS complex, and SbtA, a high-affinity low Ci inducible Na^+^/HCO_3_^−^ transporter, in WT and different mutants under conditions used for the MIMS experiments.

As expected, in WT cells grown at pH 7.5, NdhD3 and SbtA were not detected under HC conditions, but both proteins were strongly accumulated in LC (Figure 3*D*). However, in LC conditions, the increase in alkalinity of the growth medium to pH 9.0 resulted in markedly lower levels of NdhD3 and SbtA accumulation compared to those observed at pH 7.5. The effect was more pronounced in the case of SbtA. Interestingly, the Δ*flv2* and Δ*flv4* mutants demonstrated the decrease of SbtA accumulation compared to WT even at pH 7.5 in LC, whereas in *flv4/OE* SbtA remained at the same level as in WT (Figure 3D).

Thus, the expression of the SbtA protein closely followed the alterations in the expression of Flv2 and Flv4 proteins under all growth conditions, suggesting that Flv2/Flv4 and the Ci uptake mechanisms, particularly the inducible high-affinity Na^+^/HCO_3_^−^ transporter, share a common regulatory pathway of protein expression.

Unlike the growth media at pH 6 – 8.2, the Ci-pool at pH 9 contains an additional species, CO_3_^2−^. It is possible that a small amount of CO_3_^2−^ in the external growth medium acts as a signal to trigger the regulation of *flv2* and *flv4* expression via antisense RNA *as1-flv4* and the master transcription factors, *ndhR* or *cmpR* (Eisenhut et al., 2012). Considering that the double negative charge of CO_3_^2−^ prevents its diffusion through the cell membrane, and an active carbonate uptake transporter remains at present unknown, we cannot yet consider CO_3_^2−^ as an internal sensor. To get further insights into the carbonate effect on O_2_ photoreduction, the MIMS experiments were monitored on FDP mutants grown in the BG-11 medium in the presence and absence of sodium carbonate.

#### The effect of sodium carbonate

In order to control the available C_i_ species, the experimental cultures were grown in BG-11 media (pH 7.5) without sodium carbonate (Na_2_CO_3_). Culturing the cells without Na_2_CO_3_ at pH 7.5 clearly enhanced O_2_ photoreduction in the WT and all studied FDP mutants (Figure 2, middle panel). Despite such a clear variation in O_2_ photoreduction rates in WT, no significant difference in gene transcript (Figure 3B) and protein levels (Figure 3C) of FDPs were observed in the presence or absence of Na_2_CO_3_.

The following source data and figure supplement are available for Figure 3:

**Source data 1.** Transcript abundance of *flv1, flv2, flv3* and *flv4* genes

**Figure supplement 1.** *O_2_ uptake in the WT, flv4-2/OE*, Δ*flv4 and* Δ*cyd mutant*.

### 3. The FDP induced O_2_ photoreduction does not occur at PSII or the PQ-pool level

In order to establish where the Flv2/Flv4 heterodimer related O_2_ photoreduction occurs in the electron transport chain we focused on the *flv4-2*/OE mutant (grown at LC, pH 7.5, without carbonate), which showed significantly high accumulation of Flv2 and Flv4 proteins and higher O_2_ photoreduction rate than the WT (Figure 1). When linear electron transport was blocked at Cytochrome *b*_6_*f* (Cyt *b*_6_*f*) level using DBMIB as an inhibitor (Draber et al., 1970; Yan et al., 2006), both the WT (Ermakova et al., 2016) and *flv4-2*/OE mutant cells demonstrated a strong light-induced O_2_ uptake (Figure 3-Figure supplement 1). As expected, in the Δ*cyd* mutant the light-induced O_2_ uptake was not detected in the presence of DBMIB (Ermakova et al., 2016). The addition of HQNO, an inhibitor of Cytochrome *bd* quinol oxidase (Cyd), to the DBMIB-treated WT and *flv4-2*/OE completely eliminated O_2_ photoreduction. These results confirmed that Cyd was solely responsible for the observed O_2_ photoreduction occurring at the PQ-pool level.

### 4. Growth phenotype of FDP deletion mutants under fluctuating light intensities

We have previously demonstrated that the Flv1/Flv3 heterodimer enables cell growth under fluctuating light, by functioning in the Mehler-like reaction as an efficient electron sink (Allahverdiyeva et al., 2013). However, the results of the current study clearly suggest an additional involvement of the Flv2/Flv4 heterodimer in the Mehler-like reaction, particularly under conditions of LC and at pH values of 8.2 or lower (Figure 1 and 2). These findings led us to more precisely examine the combined effects of the pH of the growth medium and the fluctuating growth light conditions (FL) on the growth performance of various FDP mutants. To this end, both severe (FL20/500, when 20 µmol photons m^−2^ s^−1^ background light was interrupted every 5 min by 30 s light pulse intensity of 500 µmol photons m^−2^ s^−1^) and mild (FL50/500, when 50 µmol photons m^−2^ s^−1^ background light was interrupted every 5 min by 30 s light pulse intensity of 500 µmol photons m^−2^ s^−1^) fluctuating lights at different pH values of the growth medium were applied. In line with our previous work, the Δ*flv1*/Δ*flv3* mutant (also Δ*flv3*/Δ*flv4*) failed to grow under severe (FL20/500) light fluctuations independently of the pH of the growth medium (Figure 4; Figure 4-Figure supplement 1). Differently to the severe FL20/500 condition, under mild fluctuating light (FL50/500) the Δ*flv1*/Δ*flv3* mutant demonstrated slower growth than WT at alkaline pH of 9 (Figure. 4), and at pH 8.2 (Mustila et al., 2016, Figure 4-Figure supplement 1) and similar growth to the WT at the lower pH 7.5 (Mustila et al., 2016, Figure 4-Figure supplement 1) and pH 6 (Figure 4). Importantly, the Δ*flv4* mutant grew similarly to the WT at all studied pH levels, both under mild and severe FL conditions (Figure 4). The Δ*flv2,* Δ*sll0218* and *flv4-2/OE* mutants also demonstrated similar growth to the WT under severe FL20/500 at pH 7.5 and 8.2 (Figure 4- Figure supplement 1). The results above strongly suggest that, in contrast to the Flv1/Flv3-originated Mehler-like reaction, the Flv2/Flv4-driven O_2_ photoreduction is not essential for the survival of cells under fluctuating light.

**Figure 4.**
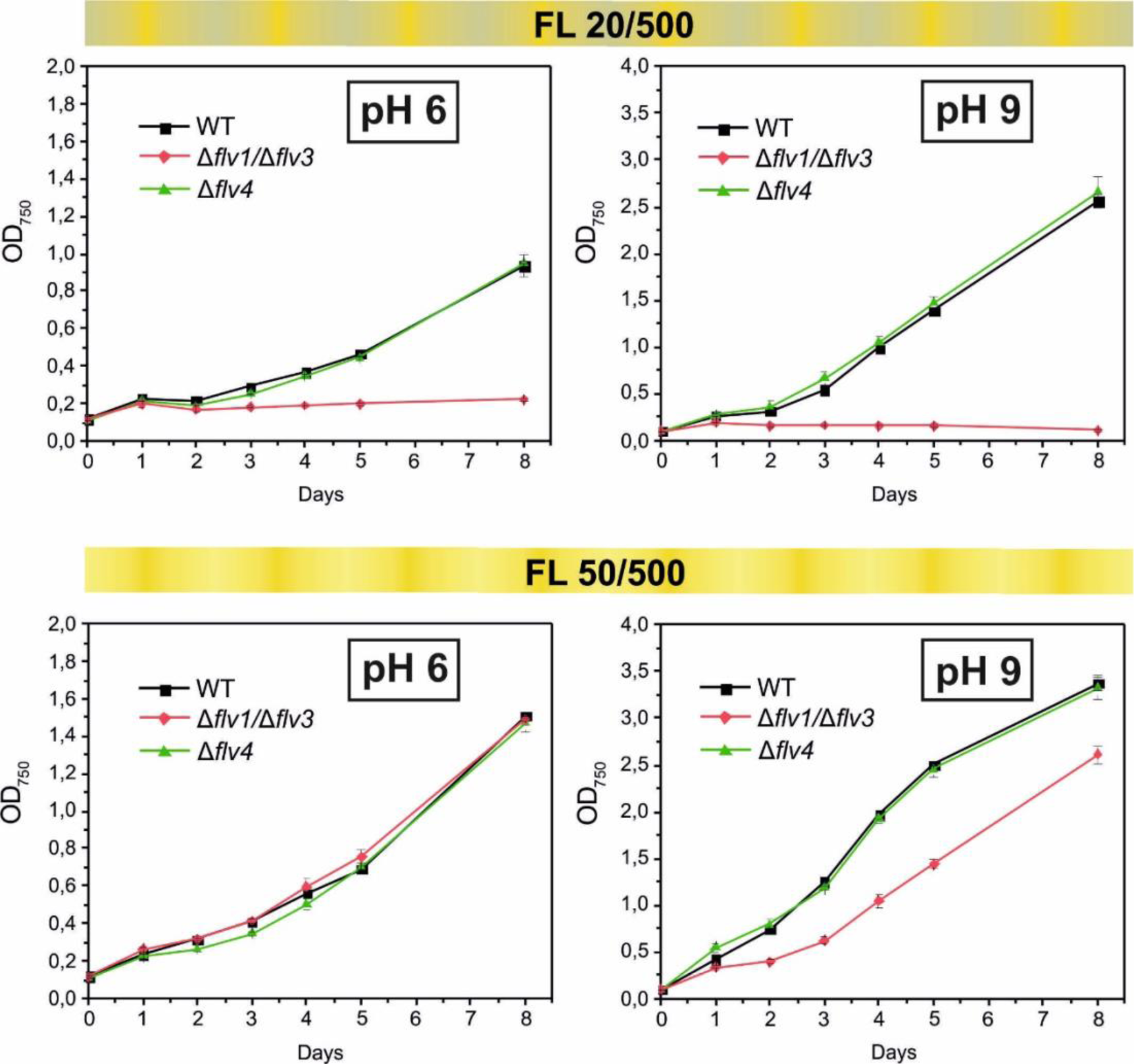
Growth curves of the different FDPs mutants under fluctuating light intensities. Pre-cultures were grown under HC for 3 days illuminated with constant light of 50 µmol photons m^−2^ s^−1^. The cells pre-grown at pH 9 or pH 8.2 (for experimental culture at pH 6) were harvested, resuspended in fresh BG-11 (pH 9 or 6), adjusted to OD_750_=0.1 and shifted to LC. Experimental cultures were grown under FL 20/500 or 50/500 regime for 8 days. The experiment was conducted in 2 independent biological replicates and average values was plotted.

The following source data and figure supplements are available for Figure 4:

**Source data 1.** Growth of the different FDPs mutants under fluctuating light intensities

**Figure supplement 1.** *Growth curves of the different Flv mutants under fluctuating light intensities*

### 5. Effect of increasing light intensities on the Mehler-like reaction

In order to assess the response of the Mehler-like reaction to different light intensities, the WT, Δ*flv4* and Δ*flv1*/Δ*flv3* mutant cells were illuminated with 500, 1000 and 1500 *µ*mol photons m^−2^ s^−1^ white light (Figure 5). Increasing light intensity from 500 to 1000 *µ*mol photons m^−2^ s^−1^ resulted in two-fold enhancement of the maximum O_2_ photoreduction rate in WT (Figure 5A, 5D). Further increase in light intensity (1500 *µ*mol photons m^−2^ s^−1^) enhanced only slightly (up to 2.3-fold) the maximum O_2_ photoreduction rate, suggesting that the applied light intensity was already saturating. Likewise, the Δ*flv4* mutant demonstrated about 1.9- and 2.3-fold enhancement in the maximum rate of transient light-induced O_2_ reduction under 1000 and 1500 *µ*mol photons m^−2^ s^−1^ illumination, respectively (Figure 5C, 5D). On the contrary, the Δ*flv1*/Δ*flv3* mutant, different from WT and Δ*flv4*, showed slower response to increasing light intensities (1.6- and 1.8-fold enhancement in the maximum rate at 1000 and 1500 *µ*mol photons m^−2^ s^−1^, respectively) (Figure 5B, 5D). It is important to note that both the Δ*flv4* and Δ*flv1*/Δ*flv3* mutants accumulate nearly the WT level of the Flv3 or Flv4/Flv2 proteins, respectively (Zhang et al., 2009; Mustila et al., 2016).

**Figure 5.**
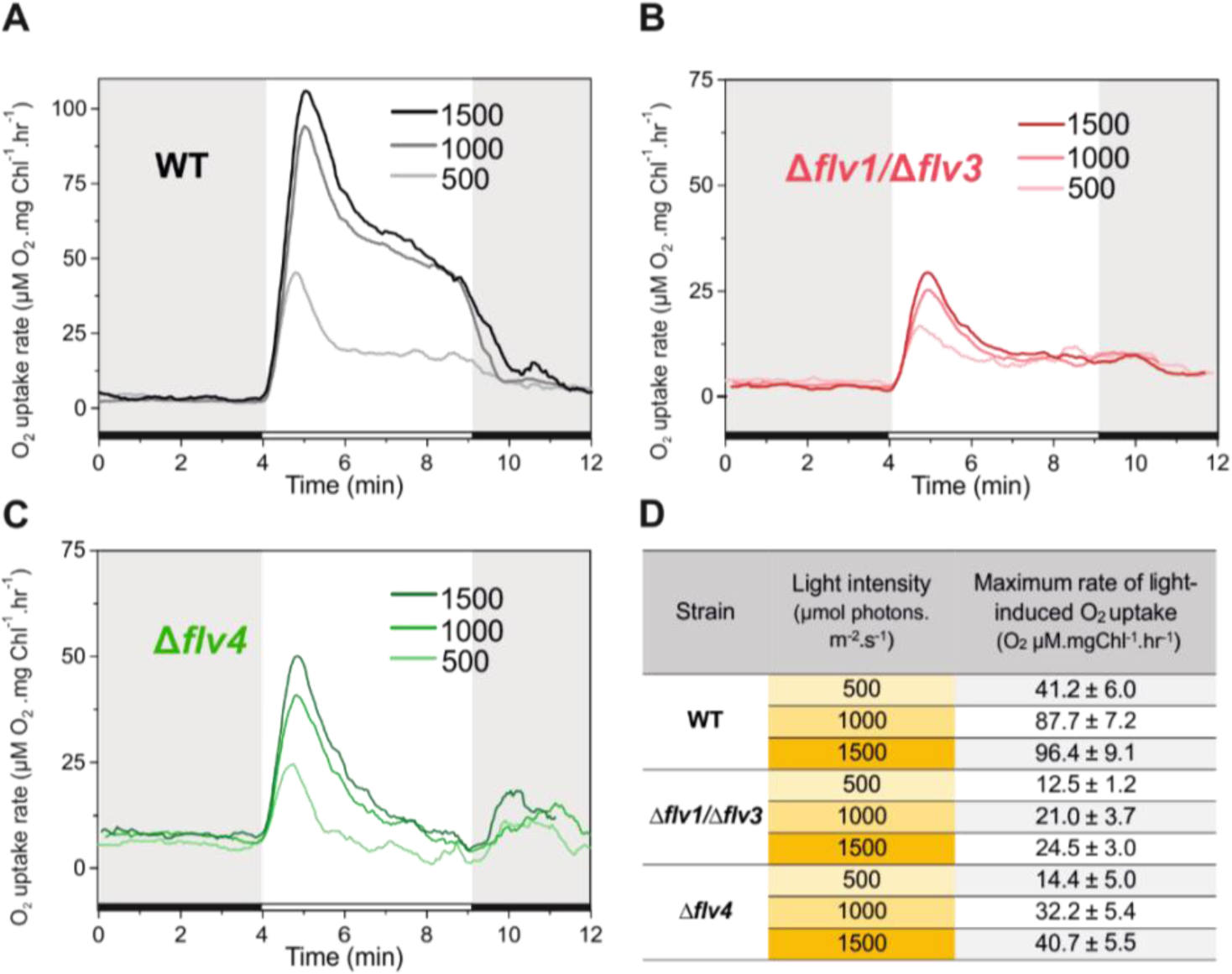
Rates of O_2_ reduction in response to increasing light intensity in WT, Δ*flv1*/Δ*flv3* and Δ*flv4* mutant cells (A, B, C, respectively). O_2_ reduction rate was recorded in darkness (grey background) and under illumination with actinic white light intensities of 500, 1000 and 1500 µmol photons m^−2^ s^−1^ (white background). Pre-cultures were grown under 3% CO_2_ (HC) at pH 7.5 for 3 days and then shifted to LC (atmospheric 0.04% CO_2_ in air) at OD750=0.2 and pH 7.5 for 4 days. For MIMS measurements, cells were harvested and resuspended in fresh BG11 medium at a Chl *a* concentration of 10 µg mL^−1^. (D) Maximum rate of light-induced O_2_ uptake (O_2_ µM. mgChl^−1^.hr^−1^) of WT, Δ*flv1*/Δ*flv3* and Δ*flv4* mutant cells at different light intensities applied. The experiment was conducted in 3 independent biological replicates and a representative plot is shown (Figure 5-source data 1).

The following source data is available for Figure 5:

**Source data 1.** Rates of O_2_ reduction in response to increasing light intensity in WT, Δ*flv1*/Δ*flv3* and Δ*flv4* mutant cells.

The fast and transient response of the Δ*flv4* mutant cells to drastic increase in the light intensity (Figure 5C) confirmed a high capacity of Flv1/Flv3-related O_2_ photoreduction as an electron sink. These results explain the essential role of Flv1/Flv3, unlike Flv2/Flv4, for the survival of cells under fluctuating light intensities. Intriguingly, both the fast induction phase {I}) and quasi-stable phase {III} of O_2_ photoreduction rate in WT were notably greater than summation of the individual O_2_ photoreduction rates from Δ*flv1*/Δ*flv3* and Δ*flv4,* thus implying a clear enhancement of O_2_ photoreduction by various oligomer activities in the presence of all four FDPs.

### 6. Growth history of cells has long-term consequences on the Mehler-like reaction

The inoculum size (starting OD_750_ value) determines how well the light penetrates into the culture upon starting the cultivation. In previous studies, cells were pre-grown in HC, then harvested at a late logarithmic phase and inoculated in fresh BG-11 (pH 8.2) at OD_750_≈0.4-0.5, before the shift to LC for the next 3 days (Allahverdiyeva et al., 2011, 2013; Ermakova et al., 2016). To ensure better light penetration to the cultures and to improve the acclimation of cells to the conditions used in this study, the experimental WT and Δ*flv1*/Δ*flv3* cultures were inoculated at a low OD_750_≈0.1-0.2 and then cultivated for 4 days (instead of 3 days in previous studies). The WT cells grown under LC from a lower OD (OD_750_≈0.2) demonstrated notably higher O_2_ uptake during illumination, compared to the cells shifted to LC at OD_750_≈0.5 (Figure. 6A). Importantly, the Δ*flv1*/Δ*flv3* mutant cells shifted to LC at a lower OD (OD_750_≈0.2) also demonstrated a residual steady-state O_2_ photoreduction activity.

**Figure 6.**
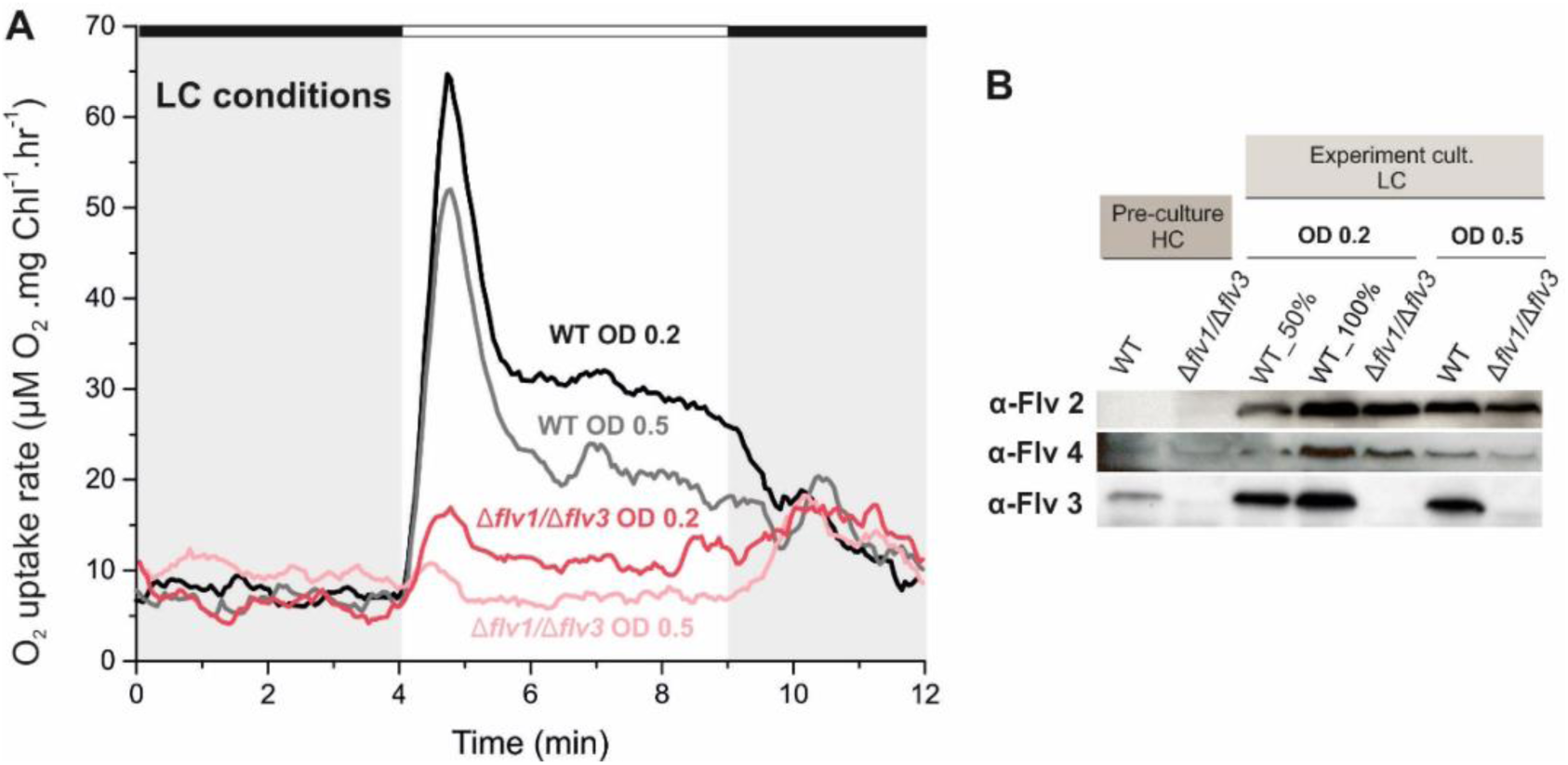
Effect of inoculum size on the O_2_ photoreduction and accumulation of FDPs in the WT and Δ*flv1*/Δ*flv3* mutant cells. (*A*) Rates of O_2_ uptake measured by MIMS during darkness (grey background) and under illumination with actinic white light at an intensity of 500 µmol photos m^−2^ s^−1^ (white background). (*B*) Protein immunoblots showing the relative accumulation of different FDPs in the WT and Δ*flv1*/Δ*flv3* mutant cells. Pre-cultures were grown in BG-11 (pH 8.2) under HC until late logarithmic phase (OD_750_≈2.5), then harvested and inoculated in fresh BG-11 under LC at OD_750_=0.2 for 4 days or OD_750_=0.5 for 3 days. The experiment was conducted in 3 independent biological replicates and a representative plot is shown in (A).

The following source data is available for Figure 6:

**Source data 1.** Rates of O_2_ reduction of WT, Δ*flv1*/Δ*flv3* and Δ*flv4* mutant cells grown at different inoculum size.

Immunoblot analysis using specific FDP antibodies showed that the WT cells transferred from HC to LC at OD_750_=0.2 accumulated higher amount of the Flv2, Flv3 and Flv4 proteins compared to the cells shifted to LC at OD_750_=0.5 (Figure 6B). A similar trend was also observed in the Δ*flv1*/Δ*flv3* mutant, which accumulated more Flv2 and Flv4 when cultivated at LC from OD_750_=0.2. This is in line with previous results showing that the accumulation of *flv2* and *flv4* transcripts in *Synechocystis* (upon a shift from HC to LC, Zhang et al., 2009) and vegetative cell-specific *flv1A* and *flv3A* transcripts in *Anabaena* sp. PCC 7120 (upon a shift from dark to light, Ermakova et al., 2013) strongly depended on light intensity.

The results above highlight the long-lasting “memory” of cells for accumulation and activity of FDPs upon a shift of cells from pre-culture conditions to different experimental conditions for assessing the specific roles of the FDPs in O_2_ photoreduction.

## Discussion

### 1. The Flv2/Flv4 heterodimer contributes to the Mehler-like reaction when naturally expressed under LC conditions or artificially overexpressed under HC

By characterizing *Synechocystis* mutants specifically affected in the accumulation of various FDPs, we show here that Flv2 and Flv4, together with Flv1 and Flv3 proteins, are involved in O_2_ photoreduction *in vivo*. Until recently, it has generally been accepted that the Flv1/Flv3 proteins safeguard PSI under both HC and LC conditions (Allahverdiyeva et al., 2013), whereas proteins, encoded by the *flv4-2* operon and highly expressed under LC, functioning in the photoprotection of PSII by, presumably, directing excess electrons from PSII to an as yet unknown acceptor (Zhang et al., 2009., Zhang et al., 2012; Shimakawa et al., 2015). The possibility of the Flv2/Flv4 contribution to O_2_ photoreduction *in vivo* was neglected due to lack of evidence for light-induced O_2_ uptake in Δ*flv1* and/or Δ*flv3* mutants (Helman et al., 2003; Allahverdiyeva et al., 2011; Allahverdiyeva et al., 2013). Thus, Flv1 and Flv3 were assumed to be solely responsible for the Mehler-like reaction. Recently, it was demonstrated that *Synechocystis* Flv4 protein expressed in *E. coli* is capable of NADH-dependent O_2_-reduction *in vitro* (Shimakawa et al., 2015). However, the reaction rate was extremely low (almost residual) compared to the activity of FDP from anaerobic bacteria (Di Matteo et al., 2008) and the enzyme showed no affinity to NADPH. Similar scenario was previously presented for the Flv3 protein, where *in vitro* studies performed on recombinant *Synechocystis* protein led to a claim that Flv3 functions as a homodimer in NADH-dependent O_2_ reduction (very low affinity to NADPH) (Vicente et al., 2002), whilst subsequent study with Δ*flv1*-OE*flv3* (or Δ*flv3*-OE*flv1*) mutants clearly demonstrated that homooligomers of Flv3 (or Flv1) do not function in O_2_ photoreduction *in vivo* (Mustila et al., 2016). Such discrepancies between the *in vitro* and *in vivo* results suggest that the *in vitro* assays conducted thus far have apparently failed to take into full consideration all the complex intracellular interactions, *e*.*g*. the involvement of Fed or FNR as an electron donor for FDPs, or the *in vitro* experiments do not necessarily demonstrate the processes occurring *in vivo*.

In this study, we provide compelling evidence for the *in vivo* contribution of Flv2/Flv4 to the O_2_ photoreduction by applying ^18^O-labelled-oxygen and real-time gas-exchange measurements to distinct FDP deletion mutants. The inactivation of *flv2* or *flv4* is shown to result in a substantial decrease of O_2_ photoreduction in the mutants compared to WT, while the overexpression of the *flv4-2* operon increases the rate of O_2_ photoreduction approximately two-fold. In addition, the possibility that the small protein Sll0218 contributes to the Mehler-like reaction is excluded (Figure 1-Figure supplement 1, compare Figure 2-Figure supplement 2 and Figure 2).

It is noteworthy that both the Δ*flv2* (deficient in Flv2 but retaining a low amount of Flv4) and Δ*flv4* (deficient in both Flv2 and Flv4) mutants showed similar inhibition of O_2_ photoreduction rates thus supporting the function of Flv2/Flv4 as a heterodimer in the Mehler-like reaction. The existence of the Flv2/Flv4 heterodimer has been proved biochemically in *Synechocystis* (Zhang et al. 2012). Nonetheless, our data do not exclude the possibility that Flv2/Flv2 and/or Flv4/Flv4 homooligomers are involved also in processes other than O_2_ photoreduction. Such a situation occurs with the Flv1 and Flv3 proteins, which contribute as homooligomers to the photoprotection of cells under fluctuating light conditions, probably *via* an unknown electron transport and/or regulatory network (Mustila et al., 2016).

The complete elimination of light-induced O_2_ reduction in the WT cells grown at pH 8.2 (Ermakova et al., 2016) or at pH 7.5 (Figure 3-Figure supplement 1.) in the presence of electron-transport inhibitors DBMIB and HQNO, blocking photosynthetic electron transport at the level of Cyt*b*_6_*f* and Cyd, respectively, suggests that no FDP-driven O_2_ photoreduction (neither by Flv1/Flv3 nor by Flv2/Flv4) does occur at the PSII or PQ-pool level. This conclusion is also supported by the fact that different from WT and the mutants deficient in FDPs, the Δ*cyd* mutant does not exhibit a light induced O_2_ uptake in the presence of DBMIB (Ermakova et al., 2016; Figure 3-Figure supplement 1).

The results discussed above clearly indicate that both the Flv1/Flv3 and Flv2/Flv4 heterodimers have a capacity to drive the Mehler-like reaction, functioning downstream of PSI.

### 2. The Flv1/Flv3 heterodimer drives a strong and steady-state O_2_ photoreduction under HC

It is generally accepted that under LC conditions, the slowing down of the Calvin-Benson cycle leads to a build-up of reduced stromal components (Cooley and Vermaas, 2001; Holland et al., 2015), which would stimulate the Mehler reaction to dissipate excess electrons (Ort and Baker, 2002), whilst in HC conditions relatively low electron flux to O_2_ via the Mehler reaction is expected. In this study we provide evidence that the HC-grown WT cells are capable of equally high O_2_ photoreduction as the respective LC-grown WT cells and of maintaining the steady-state activity at least during the first 5-10 min of illumination (Figure 1A). Compared to WT, a drastically lower O_2_ photoreduction rate in the *Δflv1*/Δ*flv3* and *Δflv3*/Δ*flv4* mutants grown in HC confirms that O_2_ uptake in these conditions is mostly due to Flv1/Flv3-driven Mehler-like reaction (Figure 1-Figure supplement 1).

It is important to note, that the O_2_ photoreduction capacity of *Synechocystis* in general correlates with the abundance of the FDPs (Figure 1 and 6). However, the protein abundance is not the only factor that determines the O_2_ photoreduction capacity. Indeed, despite the strong and steady-state O_2_ photoreduction, the HC-grown cells demonstrate nearly undetectable Flv2 and Flv4 and low amount of Flv3 compared to that under LC. Furthermore, the increase in O_2_ photoreduction rate (Figure 2, middle panel) by omitting sodium carbonate from the growth media at pH 7.5, did not correlate with any significant change in the transcript and protein levels of the FDPs, thus suggesting a possible redox regulation of the enzyme activity.

### 3. The Flv1/Flv3 heterodimer is a rapid, strong and transient electron sink, whereas Flv2/Flv4 supports steady-state O_2_ photoreduction under LC

The Mehler-like reaction of WT cells grown under LC at pH 6 – 8.2 exhibits a triphasic kinetics of O_2_ photoreduction originating from the activity of both Flv1/Flv3 and Flv2/Flv4 heterodimers (Figure 2). In this study, we were able to unravel the contribution of Flv1/Flv3 and Flv2/Flv4 heterodimers to the O_2_ photoreduction kinetics: Flv1/Flv3 is mainly responsible for the rapid transient phase, whereas Flv2/Flv4 is mostly contributing to the slow steady-state phase.

The almost complete absence of Flv2 and Flv4 proteins in WT cells grown under LC at pH 9 provides an excellent model system, where the Mehler-like reaction is naturally driven solely by the Flv1/Flv3 heterodimer, as it is the case also under HC conditions. However, in contrast to HC-grown cells, where Flv1/Flv3 can drive a steady-state O_2_ photoreduction, the cells grown under LC at pH 9 demonstrate strong but only transient O_2_ photoreduction, which decays during the first 1-2 minutes of illumination (Figure 2). The identical O_2_ photoreduction kinetics of the WT cells grown at pH 9 (accumulating Flv3 but lacking both the Flv2 and Flv4 proteins) and the Δ*flv4* mutant (accumulating Flv3 but lacking Flv4 and also Flv2), together with the complete absence of O_2_ photoreduction in the Δ*flv3*/Δ*flv4* mutant demonstrate that under LC, the Flv1/Flv3 heterodimer contributes to the Mehler-like reaction in a fast and transient manner (Figure 2). Similar conclusion was previously suggested in *Synechocystis* (Allahverdiyeva et al., 2013) and for the FlvA and FlvB proteins in *Physcomitrella patens* (Gerotto et al., 2016) and *Chlamydomonas reinhardtii* (Chaux et al., 2017; Jokel et al., 2018).

The sole contribution of Flv2/Flv4 to the Mehler-like reaction is clearly demonstrated as a steady-state O_2_ photoreduction by the Δ*flv1*/Δ*flv3* mutant grown at LC pH 6 (Figure 2), whilst the same mutant cells grown at pH 7.5 and 8.2 show only residual steady-state O_2_ photoreduction. It is important to note that the Flv2/Flv4 heterodimer, when expressed, can readily contribute to O_2_ photoreduction under HC as demonstrated by the *flv4-2*/OE strain (Figure 1), thus excluding all redox and structural hindrances for Flv2/Flv4 to function in O_2_ photoreduction at HC. However, such contribution is naturally abolished in WT cells grown under high levels of CO_2_ by down-regulation of the *flv4-2* operon (Zhang et al., 2009; Zhang et al., 2012).

The rate of the Mehler-like reaction in WT cells exceeds the cumulative O_2_ photoreduction driven solely by Flv1/Flv3 (observed in Δ*flv4*) and Flv2/Flv4 (observed in Δ*flv1*/Δ*flv3*). This demonstrates that all the four FDPs are required for an efficient Mehler-like reaction in WT cells upon growth under LC (except at pH 9). A complex interaction between FDPs possibly arises from a coordinated inter-regulation of Flv1/Flv3 and Flv2/Flv4 heterodimers and on possible occurrence of some active Flv1-4 oligomer (Figure 7). Despite detection of homotetrameric organization of *Synechocystis* Flv3 *in vitro* (Mustila et al. 2016), the direct biochemical demonstration of homo- or heterotetramers *in vivo* have not been successful.

**Figure 7.**
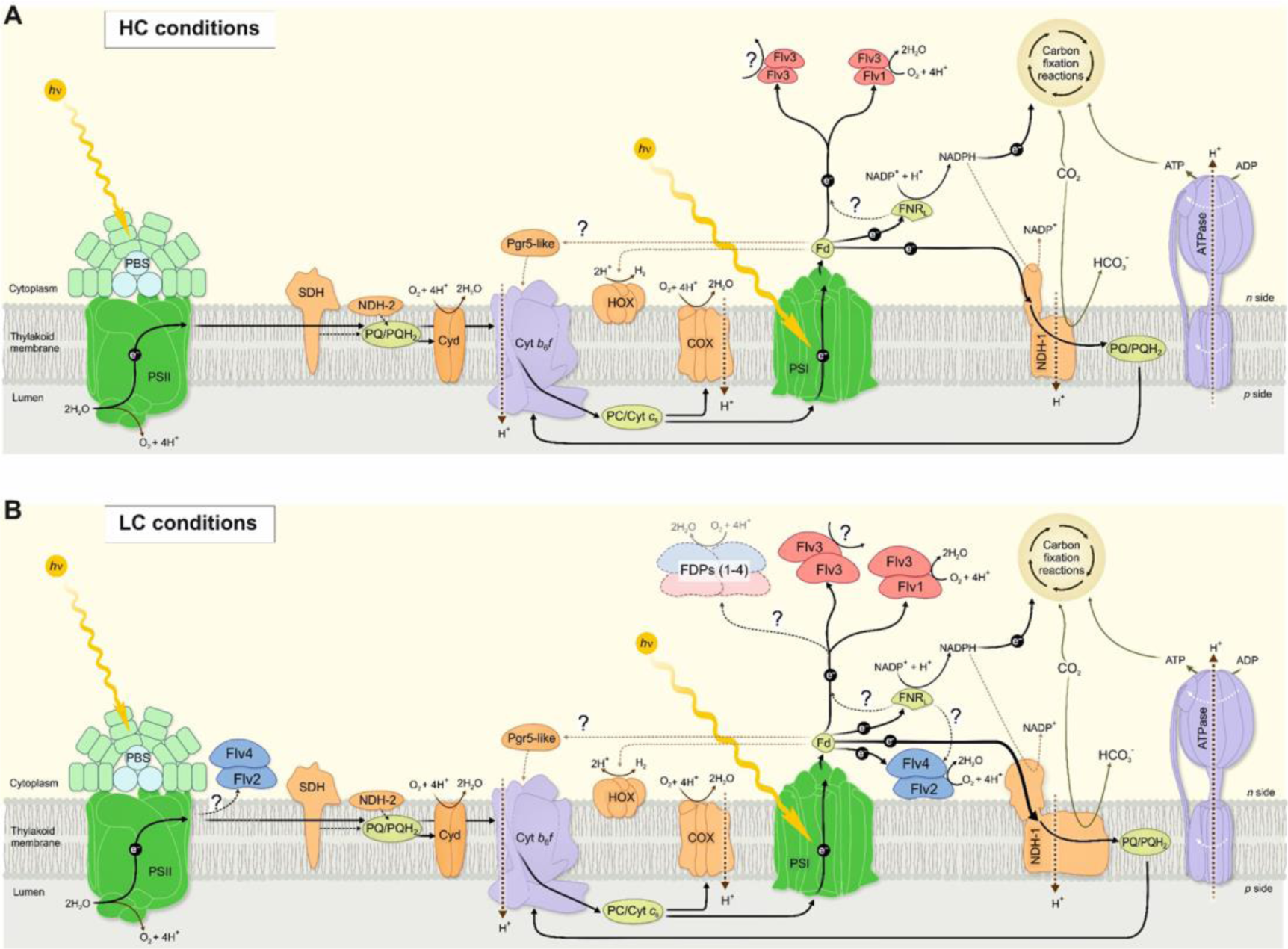
A schematic drawing of photosynthetic light reactions and alternative electron transport routes. (A) A steady-state Mehler-like reaction in HC is carried out by the low-abundant, yet catalytically efficient Flv1/Flv3 heterodimer. The Flv3/Flv3 homooligomer is involved in photoprotection as an electron valve with unknown acceptor or as a component of a signaling/regulating network (Mustila et al., 2016). (B) In LC-grown cells the two pairs of FDP heterodimers are involved in Mehler-like reaction: Flv1/Flv3 mainly drives rapid and transient O_2_ photoreduction and Flv2/Flv4 operates relatively slowly and provides a steady-state background O_2_ photoreduction. The soluble Flv1/Flv3 heterodimers function as an immediate acceptor of electrons originated from the thylakoid membrane, whereas association of Flv2/Flv4 with the thylakoid membrane (and/or Flv1/Flv3) is controlled by *pmf* and Mg^2+^. Several oligomeric forms of FDPs are hypothesized to exist, including a heterotetramer comprising different FDP protein compositions. The higher abundance of total NDH-1 complexes and FDPs oligomers in LC conditions, compared to HC conditions, is represented by larger size of the protein complexes.

The growth inhibition of Δ*flv1*/Δ*flv3* cells under severe fluctuating conditions (FL 20/500) at pH 8.2 (Allahverdiyeva et al., 2013), pH 7.5 (Mustila et al., 2016), pH 6 and pH 9 (Figure 5) demonstrate the essential role of Flv1 and/or Flv3 during drastic changes of light intensity, whereas Flv2 and Flv4 are dispensable under the same conditions (Figure 4, Figure 4-Figure supplement 1). Here, we demonstrate that the crucial importance of Flv1/Flv3 heterodimers relies on their high capacity to rapidly and effectively respond to increasing light intensities (Figure 5). By adjusting their O_2_ photoreduction activity, the Flv1/Flv3 heterodimer works as an efficient and fast sink of electrons, whereas the responsiveness of Flv2/Flv4 is relatively limited and the heterodimer mostly functions at slow time-scale in a steady-state O_2_ photoreduction.

The intracellular location of these enzymes may partially contribute to the difference in O_2_ photoreduction: Flv1 and Flv3 are soluble cytosolic proteins able to quickly associate with soluble Fed and direct electrons towards O_2_ photoreduction. In line with this, the possible interaction between *Synechocystis* Flv1, Flv3 and Fed (Hanke et al., 2011), Flv3 and Fed9 (Cassier-Chauvat and Chauvat, 2014), *Chlamydomonas reinhardtii* FLVB and FED1 (Peden et al., 2013) have been reported. The Flv2/Flv4 heterodimer, specific for cyanobacteria, was suggested to bind to the thylakoid membrane upon increase in Mg^2+^ concentration on the cytoplasmic surface of the thylakoid membrane when the lights are turned on (Zhang et al., 2012). Likely, the association of Flv2/Flv4 with the membrane enhances the electron transfer from Fed (or FNR) to Flv2/Flv4 and would probably result in a delayed and limited O_2_ photoreduction activity by Flv2/Flv4. Yet, the possibility that FDPs accept electrons from different and specific Fed paralogs cannot be excluded.

### 4. Traffic downstream of PSI affects the FDP-mediated Mehler-like reaction

The questions arise as to why Flv1/Flv3 driven O_2_ photoreduction is only transient in cells grown under LC and why an extra Flv2/Flv4 pair is required to drive the Mehler-like reaction in *Synechocystis*.

Unlike WT cells demonstrating biphasic decay kinetics of O_2_ photoreduction under LC conditions (Figure 1A and Figure 2), the M55 mutant (deficient in CET, CO_2_ uptake and respiration) (Ohkawa et al., 2000) shows steady-state O_2_ photoreduction, similar to the HC-grown WT (Figure 1B and 1D). This suggests that the strongly upregulated NDH-1 complex under LC in *Synechocystis* (Zhang et al., 2004), contributes to a rapid quenching of O_2_-photoreduction (Figure 1A, phase {II}) by efficient withdrawal of electrons from reduced Fed. Under such circumstance, the low but steady-state activity of the Flv2/Flv4 heterodimer is likely to be important for keeping linear electron transport in an oxidized state. This would explain why the PQ-pool is more oxidized in the presence of Flv2/Flv4 and more reduced in its absence, indirectly affecting PSII activity (Zhang et al., 2012; Bersanini et al., 2014 and Chukhutsina et al., 2015). Thus, by allocating the roles of FDPs between the two pairs of heterodimers (Flv1/Flv3 and Flv2/Flv4), the cells are well positioned to respond appropriately to changing C_i_ levels as well as to abrupt changes in light intensity, in a coordinated and energetically efficient manner.

Unlike prokaryotic cyanobacteria, chlorophytic algae (*e*.*g*. *Chlamydomonas reinhardtii*) and mosses rely not only on the FDP-driven pathway, but also harbor the PROTON GRADIENT REGULATION5 (PGR5)/PGR5 LIKE PHOTOSYNTHETIC PHENOTYPE 1 (PGRL1) pathway which operates concomitantly to protect the cells under fluctuating light. It is noteworthy, however, that the PGR5/PGRL1 machinery in *Chlamydomonas reinhardtii* is neither fast nor strong enough to mitigate acceptor-side pressure under highly fluctuating light intensities. To complement this deficiency, the FDP-mediated pathway is indispensable for coping with sudden increases in light intensity (Jokel et al., 2018). Interestingly, the introduction of *Physcomitrella patens* FDPs rescues a fluctuating light phenotype of the PGR5 *Arabidopsis thaliana* mutant (Yamamoto et al., 2016; 2019), and alleviates PSI photodamage in the PGR5-RNAi, *crr6* (defective in NDH-dependent CET) and the PGR5-RNAi *crr6* double mutants of *Oryza sativa* by acting as a safety valve under fluctuating light and substituting for CET without competing with CO_2_ fixation under constant light (Wada et al., 2017). Moreover, the expression of *Synechocystis* Flv1 and Flv3 in tobacco plants enhances photosynthetic efficiency during dark-light transitions by providing an additional electron sink (Gómez et al., 2018). Although data on Flv2/Flv4 proteins expressed in angiosperms is not yet available, our results collectively suggest that the FDP pathway(s) is important to consider in future high-yield crop development and microbial cell factories.

Figure 7 provides a summary scheme of our understanding of the function and interaction of the different FDPs and their oligomers in photoprotection of the photosynthetic apparatus in the model cyanobacterium *Synechocystis* sp. PCC 6803. The importance of the available Ci species in the function and accumulation of FDPs is emphasized by separate schemes for the HC and LC growth conditions.

## Materials and Methods

### Strains and culture conditions

The glucose-tolerant *Synechocystis* sp. PCC 6803 was used as wild type (WT) strain (Williams, 1988). The FDP inactivation mutants Δ*flv2*, Δ*flv4* (Zhang et al., 2012), and the double mutants Δ*flv1*/Δ*flv3* (Allahverdiyeva et al., 2011), and Δ*flv3*/Δ*flv4* (Helman et al., 2003) have been described previously. The *flv4-2*/OE and Δ*sll0218* mutants were described in (Bersanini et al., 2014; Bersanini et al., 2017).

Pre-experimental cultures were grown at 30°C in BG11 medium, illuminated with continuous white light of 50 µmol photons m^−2^ s^−1^ (growth light: GL), under air enriched with 3% CO_2_ (high carbon: HC). BG-11 medium was buffered with 20 mM 2-(N-morpholino) ethanesulfonic acid (MES, pH 6.0), 20 mM HEPES-NaOH (pH 7.5), 10 mM TES-KOH (pH 8.2) or 10 mM N-Cyclohexyl-2-aminoethanesulfonic acid (CHES, pH 9.0), according to the pH of the experimental condition. Pre-cultures were harvested at logarithmic growth phase, inoculated in fresh BG-11 medium at OD_750_=0.2 (or OD_750_ = 0.5 when mentioned), measured with and shifted to low CO_2_ (atmospheric 0.04% CO_2_ in air, LC). OD_750_ was measured using Lambda 25 UV/VIS spectrometer (PerkinElmer, USA). HC experimental cultures were inoculated at OD_750_= 0.1 and kept at HC for 3 days. During experimental cultivation, cells were grown under continuous GL at 30°C with agitation at 120 rpm and without antibiotics. For growth curves, cells pre-cultivated under continuous GL and HC were collected, inoculated at OD750= 0.1 and shifted to LC under a light regime with a background light of 20 µmol photons m^−2^ s^−1^ interrupted with 500 µmol photons m^−2^ s^−1^ for 30 s every 5 min (FL 20/500) or 50 µmol photons m^−2^ s^−1^ interrupted with 500 µmol photons m^−2^ s^−1^ for 30s every 5 min (FL 50/500). Sodium carbonate was omitted from the BG-11 ingredients when mentioned.

Absence of contamination with heterotrophic bacteria was checked by dropping liquid culture on LB and R2A agar plates and kept at 30°C.

### Isolation of total RNA and Real-time quantitative PCR (RT-qPCR)

Total RNA was isolated from exponentially growing *Synechocystis* by hot-phenol method previously described (Tyystjärvi et al., 2001). After removing any residual genomic DNA, the RNA concentration and purity were measured with a NanoDrop spectrophotometer (Thermo Scientific, USA). RNA integrity was verified by agarose gel electrophoresis.

Complementary DNA was synthesized from 1 μg of purified RNA using the iScript cDNA Synthesis Kit (BioRad, USA) according to the manufacturer’s protocol. Synthesized cDNA was diluted four-fold and used as template for the RT-qPCR. The samples for RT-qPCR were labeled by iQ SYBR Green Supermix (BioRad, USA) to detect accumulation of amplicons in 96-well plates. The primers to detect transcripts of *flv1* and *flv2* as well as for the reference genes *rnpB* and *rimM* are described in Mustila et al. 2016. The forward and reverse primers for *flv3* were 5’-CAACTCAATCCCCGCATTAC-3’ and 5’-CAGTGGAGATTCGGAGCACT-3’ and for *flv4* 5’-ACGATGCCTGGAGTCAAAAC-3’ and 5’-GGGTATCCGCCACACTTAGA-3’. The PCR protocol was as follows: 3 min initial denaturation of cDNA at 95°C, followed by 40 cycles of 95°C for 10 s, annealing in 57°C for 30 s and extension in 72°C for 35 s. A melting curve analysis was performed at the end. Relative changes in the gene expression were determined using the qbase+ software by Biogazelle. One-way ANOVA analysis performed with SigmaPlot was used to determine significant changes in gene expression.

### MIMS experiments

*In vivo* measurements of ^16^O_2_ (mass 32) and ^18^O_2_ (mass 36) exchange was performed using a Membrane-inlet mass spectrometry (MIMS) as described previously in (Mustila et al., 2016). Cells were harvested, adjusted to 10 µg Chl *a* mL^−1^ in fresh BG-11 medium and acclimated for 1 h to the same experimental conditions as was applied for the cultivation.

### Protein Isolation, electrophoresis and immunodetection

Total cell extracts and the soluble fractions of *Synechocystis* cells were isolated as described (Zhang et al., 2009). Proteins were separated by 12% (w/v) SDS-PAGE containing 6M urea and transferred onto a PVDF membrane (Immobilion-P; Millipore, Germany) and immunodetected by protein specific antibodies.

## Figure supplements and 970 source data files available

**Figure 1-Figure supplement 1.***O_2_ photoreduction rates under high CO_2_*.

**Figure 2-Figure supplement 1.***O_2_ photoreduction rates during the dark-to-light transition of WT cells with and without addition of 1.5 mM NaHCO_3_ prior MIMS measurements*.

**Figure 2-Figure supplement 2.***O_2_ photoreduction rates of the Δflv2 and Δsll0218 mutants grown at LC pH 7.5* and 8.2 *with and without Na_2_CO_3_*.

**Figure 3-Figure supplement 1.***O_2_ uptake in the WT, flv4-2/OE, Δflv4 and Δcyd mutant*.

**Figure 4-Figure supplement 1.***Growth curves of the different Flv mutants under fluctuating light intensities*.

**Figure 1-source data 1.***O_2_ reduction rates of WT, flv4-2/OE and M55 mutants grown* in different CO_2_ levels.

**Figure 2-source data 1.***O_2_ reduction rates of WT and FDP mutants grown at different pH levels*.

**Figure 3-source data 1.***Transcript abundance of flv1, flv2, flv3 and flv4 genes*.

**Figure 4-source data 1.***Growth of the different FDPs mutants under fluctuating light intensities*

**Figure 5-source data 1.***Rates of O_2_ reduction in response to increasing light intensity in WT*, Δ*flv1*/Δ*flv3 and* Δ*flv4 mutant cells*.

**Figure 6-source data 1.***Rates of O_2_ reduction of WT*, Δ*flv1*/Δ*flv3 and* Δ*flv4 mutant cells grown at different inoculum size*.

