## Supplementary file for "Flv1-4 proteins function in versatile combinations in O_2_ photoreduction in cyanobacteria"

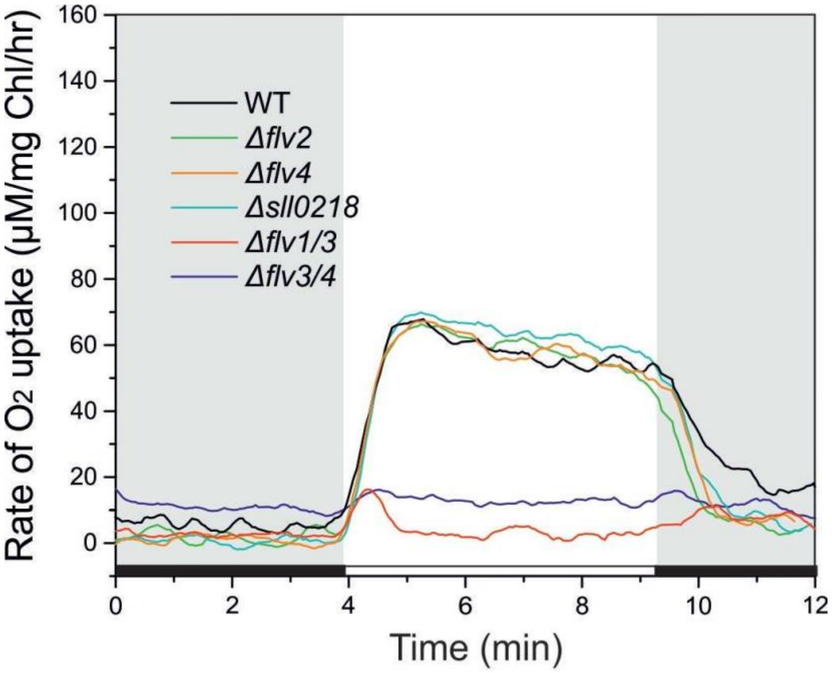


***Figure 1- Figure supplement 1*. $O_2$ photoreduction rates under high $CO_2$.** *Cells were grown under 3% $CO_2$ (BG11, pH 8.2), harvested and resuspended in fresh BG 11 at Chl 10 ug/ml. $O_2$ uptake was recorded during the transition from dark to high-light (500 µmol photons $m^{-2}$ $s^{-1}$).*

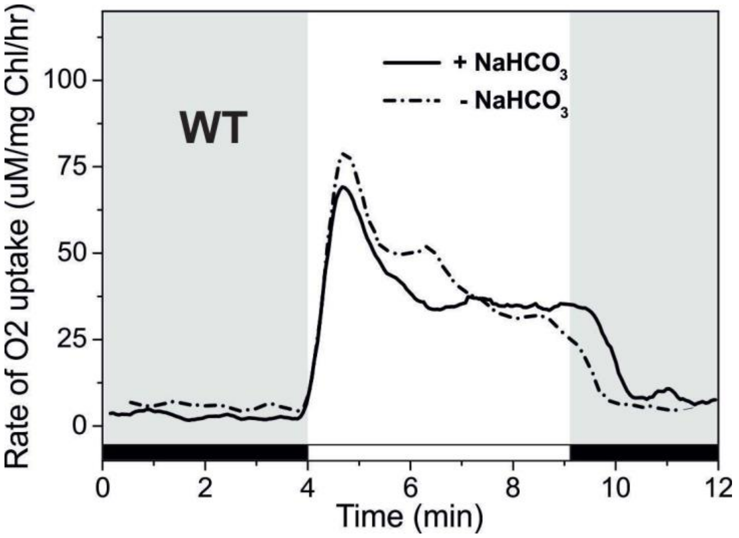


***Figure 2- Figure supplement 1. $O_2$ photoreduction rates during the dark-to-light transition of WT cells with and without addition of 1.5 mM $NaHCO_3$ prior MIMS measurements.*** *The cells were harvested and inoculated in the fresh BG11 7.5 without $Na_2CO_3$. Prior to MIMS measurement, cells were supplemented with 1.5 mM $NaHCO_3$ (solid line), or measured in the absence of an additional carbon source.*

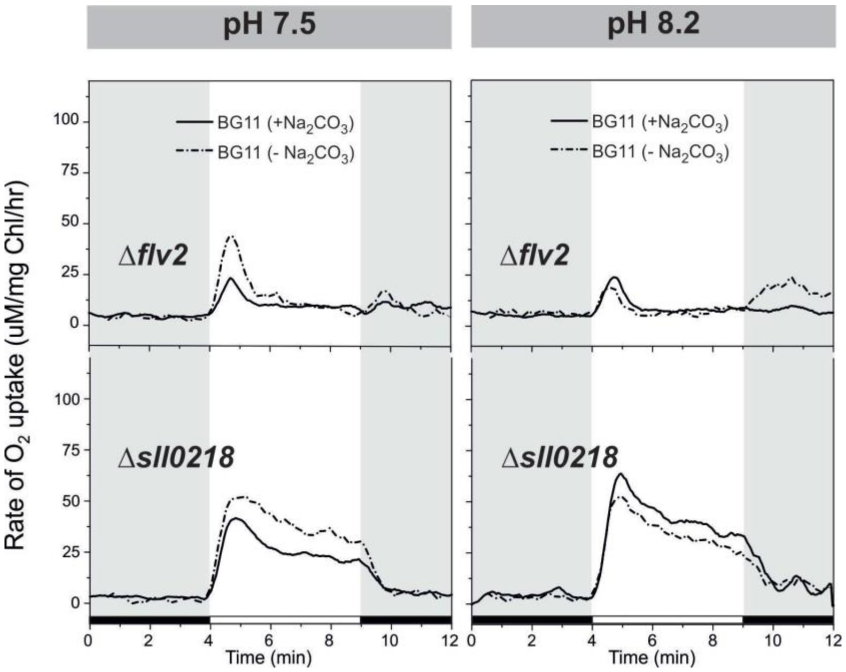


***Figure 2- Figure supplement 2. $O_2$ photoreduction rates of the Δflv2 and Δsll0218 mutants grown at LC pH 7.5* and 8.2 *with and without $Na_2CO_3$.*** *Pre-cultures were grown under HC for 3 days at pH 7.5 or pH 8.2 in BG11 media with or without $Na_2CO_3$. For $O_2$ photoreduction experiments, cells were shifted to LC at $OD_{750}$≈0.2 and grown for 4 days.*

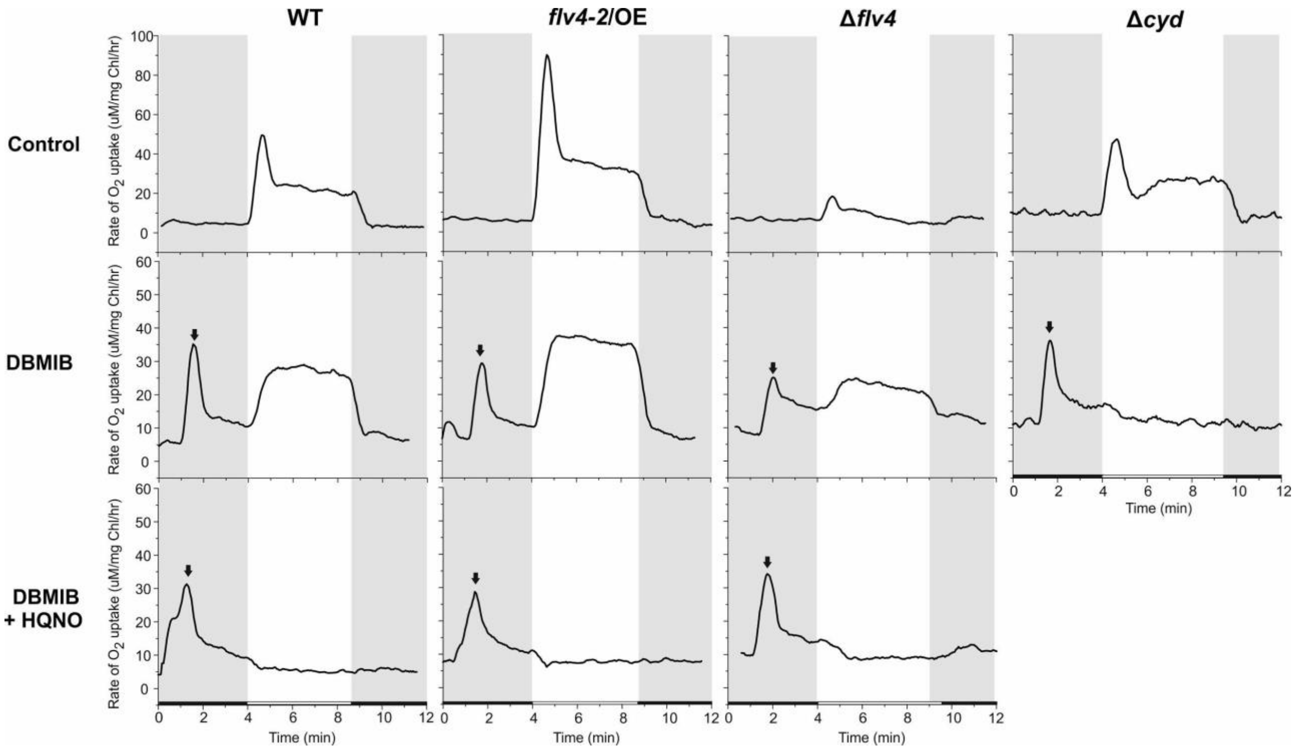


***Figure 3- Figure supplement 1. $O_2$ uptake in the WT, flv4-2/OE, Δflv4 and Δcyd mutant.*** *The cells were grown at LC in BG11 at pH 7.5. 25uM DBMIB and 50uM HQNO were added directly to the cuvette immediately prior to MIMS measurement. The arrow indicates the time when inhibitor was added to the sample. The Δcyd mutant was previously described in* Howitt and Vermaas, 1998.

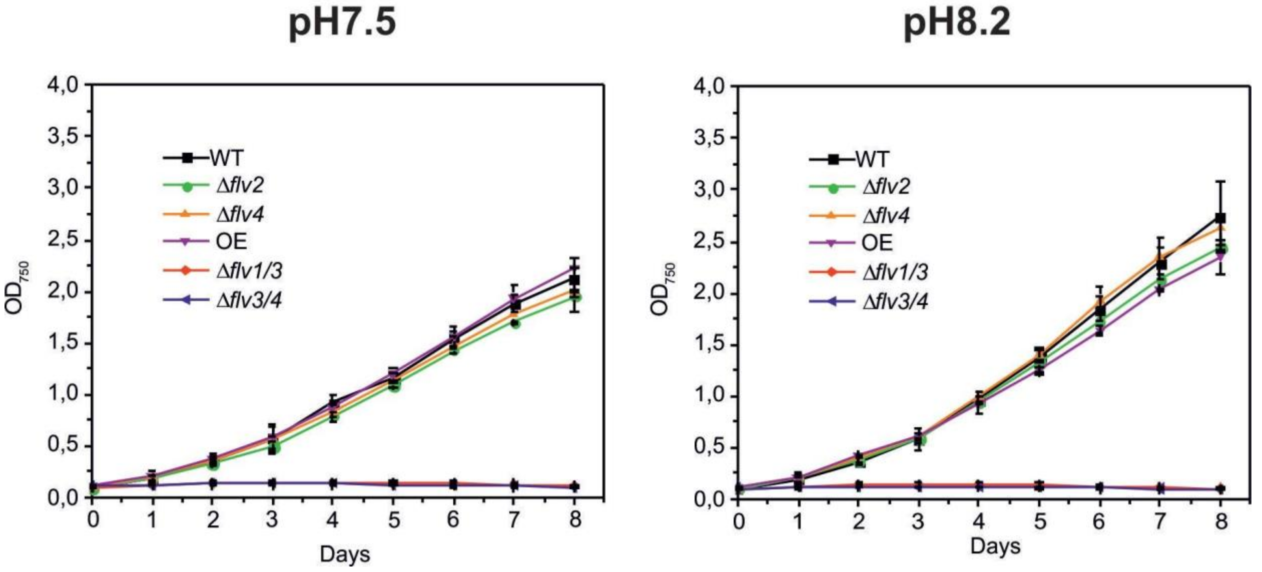


***Figure 4- Figure supplement 1. Growth curves of the different Flv mutants under fluctuating light intensities*** *(FL20/500 - 20 μmol photons $m^{-2}s^{-1}$background light is interrupted with 30 s of 500 μmol photons $m^{-2}s^{-1}$light every 5 min). Cells were grown in BG11 (pH 7.5) in the absence of $Na_2CO_3$ and shifted from HC to LC at pH 7.5 or pH 8.2*
